# A cocktail of B vitamins with nicotinamide riboside, folate and cobalamin preserves cardiac function and mitochondrial oxidative capacities in a mouse model of heart failure

**DOI:** 10.1101/2025.02.18.638803

**Authors:** Solène E. Boitard, Morgane Delouche, Ahmed Karoui, Mélanie Gressette, Iman Momken, Bertrand Bouchard, Françoise Mercier-Nomé, Apolline Imbard, Christophe Lemaire, Anne Garnier, Matthieu Ruiz, Mathias Mericskay, Jérôme Piquereau

**Author notes:** Corresponding authors: Dr. Jérôme Piquereau, Dr. Mathias Mericskay. these authors contributed equally to this work (Mathias Mericskay, Jérôme Piquereau).

## Abstract

Despite a substantial therapeutic arsenal to treat patients affected by heart failure (HF), no treatment specifically targets alterations of cardiac energy metabolism and mitochondrial functions. Yet, these alterations are now well-known and their involvement in HF pathophysiology has been demonstrated for years. Based on the results of previous studies demonstrating the cardiac preventive effects of B vitamins when introduced before inducing cardiac pressure overload in mice, we investigated the efficacy of a diet supplemented with a B vitamin cocktail (B3, B9 and B12 (3VitB)) to restore energy metabolism and improve cardiac function in an animal model of established HF. Heart Failure was induced by transverse aortic constriction (TAC) in male and female C57Bl6N mice and 3VitB treatment was introduced four weeks later in animals meeting criteria of heart failure with restricted ejection fraction. A 20-week survival study showed a significant longer life expectancy in TAC males treated with Vit3B in comparison with TAC males fed with normal diet, and this was associated with a reduction in the over time TAC-induced alterations of ejection fraction, stroke volume, and systolic and diastolic left ventricular diameter. Although, these effects on survival and cardiac function were less clear in females due to their higher resistance to TAC, the Vit3B cocktail was beneficial in females as 8 weeks of treatment improved physical capacities and led to milder cardiomyocyte stress-induced hypertrophy in similar ways to those observed in males. In both sexes, 3VitB treated TAC mice exhibited higher mitochondrial oxidative capacities than TAC mice fed with normal diet. This was at least partly supported by the maintenance of the mitochondrial biogenesis process activation, demonstrated by the higher expression of genes such as *Tfam* and NRF1 protein level in 3VitB treated TAC groups. Interestingly, our results revealed sex-specificities not only in response to cardiac pressure overload but also in response to 3VitB treatment that acted through different mechanisms that involved AMPK in males and SIRT1 in females. Overall, this study demonstrated the efficacy of 3VitB to preserved cardiac function and energy metabolism in an established HF model, especially in males that are more sensitive to cardiac pressure overload. This confers credit to vitamin supplementations and to metabolic therapy as new strategies in the treatment of HF.

## Introduction

Despite constant advances in medical science, prevalence of heart failure (HF) is still high and is expected to rise in the world with ageing of the population and the spread of western diet and associated comorbidities. This syndrome, which is often the terminal stage of diverse chronic cardiovascular diseases (CVD), is a major cause of death worldwide and cannot be fully cured in spite of important efforts to improve therapeutics these last decades.

It is now well-established that profound modulations of energy metabolism are involved in the development of HF. The failing heart is in an energy-depleted state (Neubauer, 2007), which is associated with severe mitochondrial dysfunctions and a shift from predominant fatty-acid oxidation to an increased carbohydrate use (Davila-Roman *et al*, 2002; Doenst *et al*, 2013). While the underlying reasons for such alterations remain only partially understood, the knowledge acquired from HF animal models and HF patients on the dysregulations of mitochondrial life cycle (Ventura-Clapier *et al*, 2011) and the disruption of nicotinamide adenine dinucleotide (NAD+) homeostasis (Matasic *et al*, 2018; Mericskay, 2016) confers credit to the concept of a metabolic therapy that would aim to restore the energy metabolism of the failing heart. Yet, none of the main drugs in the therapeutic arsenal currently used to treat HF in clinical practice (β-blockers, angiotensin converting enzyme 2 inhibitors (ACE) inhibitor, mineralocorticoid receptor antagonists) specifically address this issue of altered cardiac energy metabolism, thereby prompting researchers to find new molecules that could target key players of cellular energy.

In this context, our team published two studies demonstrating the potential therapeutic capacity of vitamins that open new perspectives in this field (Diguet *et al*, 2017; Piquereau *et al*, 2016). According to these works, cobalamin (B12), folate (B9) and nicotinamide riboside (NR) (B3), which are part of the large vitamin B family involved in many metabolic pathways (Piquereau *et al*, 2021), could be of great interest to protect energy metabolism and function of the heart facing hemodynamic stress. Our first study (Piquereau *et al*., 2016) stipulates that the protective effects of cobalamin and folate seems to rely on their ability to impact the homocysteine/methionine cycle that would modulate the activity of the peroxisome proliferator-activated receptor gamma co-activator 1 α (PGC-1α), a master regulator of energy metabolism and mitochondrial life cycle, by promoting its methylation by protein arginine methyltransferase 1 (PRMT1) and its deacetylation by sirtuin-1 (SIRT1), two activating post-translational modulations of this co-activator (Jeninga *et al*, 2010; Teyssier *et al*, 2005). Like other studies suggested before (Gu *et al*, 2014; Tanno *et al*, 2010), these observations place SIRT1 at the heart of a mechanism that could improve cellular energy in the frame of a therapy of CVD.

SIRT1 is a deacetylase that regulates many facets of energy metabolism and its central role makes it a prime target for the development of metabolic therapies of HF. However, its activity requires NAD^+^ (Li, 2013) and the loss of myocardial NAD^+^ in HF (Mericskay, 2016) could be a hindrance to the development of such a therapy. Indeed, this lack of NAD^+^ could limit the effect of SIRT1 activation strategies on the one hand and on the other hand fully efficient catabolism requires NAD^+^ at many stages, especially for mitochondrial functions (Chini *et al*, 2017); this implies that suitable NAD^+^ level would be a prerequisite for any effective metabolic therapy. In an interesting way, in our second aforementioned study (Diguet *et al*., 2017) our team showed that treatment with NR restores cardiac NAD^+^ level and protects cardiac function in mouse models of dilated cardiomyopathy or TAC induced-pressure overload. According to our analysis, in these models NR would take advantage of an increase in the expression of the NMRK2 enzyme that catalyzes the synthesis of nicotinamide mononucleotide (NMN) (the immediate upstream precursor of NAD+) from NR, thereby compensating for the decrease in NAMPT activity which ensures NAD^+^ production from nicotinamide (NAM) in the healthy heart (Liu *et al*, 2018). Thus, NR positions itself as a key molecule to maintain a high myocardial NAD^+^ level that could form the basis of any new strategies aiming at restoring cardiac energy metabolism of the failing heart.

Based on these encouraging results, the present study consisted in testing the effect of a vitamin cocktail containing NR, folate and cobalamin in a murine model of pressure overload-induced HF. While the effects of these three B vitamins were assessed during the development of HF in our previous studies, i.e. introduction of the treatment before the appearance of the first symptoms of HF, here we decided to assess the efficacy of the vitamin B3-9-12 cocktail in a model of established HF to reveal its potential effects at a more advanced stage of the disease, as would be the case with HF patients.

## Materials and methods

### Animals

All animal experimental procedures were approved by animal ethics committee of Paris-Sud University, authorized by French government (authorization number: APAFIS#24778-2020032416337862) and complied with directive 2010/63/EU of the European Parliament on the protection of animals used for scientific purposes.

### Pressure overload-induced Heart failure model (TAC)

The mouse model of heart failure used for this study was a pressure overload induced obtained by the surgical Transverse Aortic Constriction (TAC). Forty-nine to fifty-five days old male and female C57BL/6NCrl mice (Charles River Laboratories, breeding site Italy) were anesthetized by intraperitoneal injection (0.5-mL 29G syringe) of 100 mg/kg ketamine (Imalgene 1000, Merial, France) combined with 10 mg/kg xylazine (Rompun 2%, Bayer healthcare, France) and placed under artificial respiration (MiniVent 845 - Small Animal Ventilator, Harvard Apparatus, France). Mice were remedicated by subcutaneous injection of 1 mg/kg meloxicam (Metacam, Boehringer Ingelheim, Germany) to manage the pain of the animal during surgery. A subcutaneous injection of 4 mg/kg lidocaine (Lurocaine, Vetoquinol, France) was then given at the site of the incision. The animal was kept on a hot plate (Bioseb, France) at 37°C throughout the surgery and occular gel (Lubrithal, Centravet, France) was applied to prevent drying of the eyes. Endotracheal intubation was performed to place the mouse under artificial respiration (MiniVent 845 - Small Animal Ventilator, Harvard Apparatus, France) at the frequency of 200 strokes by min for a stroke volume of 200μL. A suprasternal skin incision was made follow by a mini proximal sternotomy on the upper thorax and the first rib was opened. The thymus was retracted and strap muscles were dissected to expose the aortic arch. A 7-0 silk suture has been passed under the aorta between the origins of the brachiocephalic and left common carotid arteries. The transverse aorta constriction (TAC) was done by a ligature around a 27-gauche needle used as a guide on the aorta. After banding, the 27-gauge needle was removed and strap muscles were stitched. Chest and skin were closed by discontinous suture point. Control mice were subjected to an identical procedure without placement of a ligature (Sham). The animals were rehydrated with physiological saline at 37°C and extubated as soon as they could breathe spontaneously. Upon awakening, each mouse was isolated in a cage in a quiet and warm place. The mice received an injection of 1 mg/kg meloxicam 24 h post-op and and every 24 h for up to 72 h. If the pain of TAC was intense, an injection of 0.04 mg/kg buprenorphine (Buprécaire, Axience SAS, Pantin, France) was given. Pain management after surgery are detailed in figure S1C and supplemental methods.

### Treatment administration on non-synthetic diet

Four weeks after surgical transverse aortic constriction (TAC), mice were randomized based on different inclusion criteria assessed by trans-thoracic echocardiography: pressure gradient above 60mmHg (Figure S2B), decrease of 10% of Left Ventricular Ejection fraction (LVEF) and increase of 30% of Left Ventricular mass (Figure S2C). Mice that did not match two or more of these criteria were excluded from the study. After the randomization step, male and female mice were divided into three groups: 1) sham-operated mice receiving standard SAFE^®^ A04 non-synthetic diet (SAFE, Augy, France), 2) TAC mice receiving standard SAFE^®^ A04 non-synthetic diet called TAC-ND, 3) TAC mice receiving customized SAFE^®^ A04 supplemented with NR (4g/kg), cobalamin (1mg/kg) and folate (20 mg/kg) called TAC-3VitB in this manuscript. Animals were housed under temperature-controlled conditions (21°C) and had free access to water and to their respective food. Supplementation with vitamins started four weeks after TAC and two different treatment times were evaluated in two studies (Figure S1A and B): 1) 20 weeks of vitamins treatment for survival and functional study on males (Sham n=11; TAC-ND n=20; TAC-Vit n=19) and female (Sham n=11; TAC-ND n=16; TAC-Vit n=20); 2) 8 weeks of vitamins treatment for studying physical capacities, metabolomic, histologic, and molecular analysis on males (Sham n=9; TAC-ND n=11; TAC-Vit n=11) and females (Sham n=6; TAC-ND; n=10; TAC-Vit n=10).

### Survival

A survival study was carried out during 20 weeks of treatment to evaluate the impact of treatment on lifetime on males and females. A regular monitoring of mice body weight (each week) and cardiac function by trans-thoracic echocardiography (every four weeks) were done and correlated to end points determined on ethical project. When mice lost more than 20% of total body weight, a left ventricular ejection fraction under 20%, and end points have been reached, mice were euthanized and marked on survival curve as an event.

### Effort test on treadmill

Before introduction and after 8 weeks of treatment, the heart’s ability to adapt to physical effort was evaluated by an exertion test on treadmill (Ugo Basile, France) on male and female mice. Three days before the effort test, an adaptation phase was done with the following program: three and two days before the test, the mice were subjected to the following training session during which the treadmill started at a 3 m.min^-1^ speed that was then increased by 1 m.min^-1^ every 2 min for 10 min without slope. After a day off, the exhaustion test was done with the following program: after an acclimation of 30 min, the treadmill started at a 3 m.min^-1^ speed that was increased by 3 m.min^-1^ every 3 min without slope to determine the maximal running time on increasing speed.

### Echocardiography

Every four weeks, cardiac function was evaluated by trans-thoracic echocardiography with a Vevo 3100 device (FUJIFILM Visualsonics Inc., Toronto, Canada) supplied with a linear array 22–55 MHz MicroScan mouse cardiovascular transducer (MS550. Echocardiographic measures were performed on mice anesthetized using isoflurane (Centravet, Plancoet, France), 3% for induction and 1 to 2% for the maintenance of anesthesia. Heart Rate, body temperature, and ECG were monitored by sensors on the echography warm plate. Systolic cardiac function parameters such as left ventricular ejection fraction (LVEF), left ventricular shortening fraction (LVSF), left ventricular mass (LV mass), cardiac diameters (left ventricular end-systolic diameter (LVESD) and left ventricular end-diastolic diameter (LVEDD) and volumes (left ventricular end-systolic volume (LVESV) and left ventricular end-diastolic volume (LVEDV), and stroke volume (SV) were obtained by anatomic Bidimentional and M-mode views in parasternal long axis and parasternal short axis. Diastolic cardiac function parameters such as E/A ratio were obtained by apical four-chamber view. The distal transverse aortic flow velocity (distal to constriction in TAC mice) was measured by pulsed wave (PW) Doppler to evaluate the TAC-induced pressure gradient using the modified Bernoulli equation (Pressure gradient = 4*Velocity2). Images were analyzed by VevoLab Software (FUJIFILM Visualsonics Inc., Toronto, Canada).

### Histological analysis

Hearts were fixed in 4% paraformaldehyde, paraffin embedded and serially sectioned (3 μm). Sections were stained with Sirius red (Ab150681; Abcam) for studying fibrosis or with FITC-labeled wheat germ agglutinin (WGA) to label cell membranes. The slides were scanned using NanoZoomer2.0-RS^®^ (Hamamatsu Photonics) digital scanner. Using Image J software, fibrosis and cross-sectional area of the fibers were quantified from 12 to 15 fields of a transversal section of the left ventricle for each animal.

### Mitochondrial oxidative capacities assays

Mitochondrial respiration was studied *in situ* in saponin-permeabilized cardiac muscle fibers using a Clarke electrode (O2k-Fluorespirometer, Oroboros Instruments) as previously described (Kuznetsov *et al*, 2008). Briefly, fibers were dissected from the free wall of freshly harvested left ventricles in S solution (CaK_2_ ethyleneglycol tetraacetic acid (EGTA) (2.77mM), K_2_EGTA [100 nM free Ca2+] (7.23 mM), MgCl_2_ [1 mM free Mg2+] (6.56 mM), Na_2_ATP (5.7 mM), phosphocreatine (15 mM), taurine (20 mM), dithiothreitol (DTT) (0.5 mM), K-methane sulfonate (50 mM), imidazole (20mM), pH 7.1) on ice. Fibers were permeabilized in S solution added with saponin (50µg/ml) during 30 min at 4°C. About 5 mg (wet weight) of permeabilized fibers were placed in each oxygraphic chamber containing respiration solution (CaK_2_ ethyleneglycol tetraacetic acid (EGTA) (2.77mM), K_2_EGTA (7.23 mM), MgCl_2_ (1.38 mM), K_2_HPO_4_ (3mM), taurine (20 mM), dithiothreitol (DTT) (0.5 mM), K-methane sulfonate (90 mM), Na-methane sulfonate (10 mM), imidazole (20mM), 2 mg.ml^-1^ bovine serum albumin, pH 7.1) the temperature of which is monitored at 23 °C. Oxygen consumption of the fibers were measured after successive addition of pyruvate 1 mM, malate (4 mM), ADP (2 mM), succinate (15 mM), amytal (an inhibitor of complex I, 1 mM) and N,N, N’,N’-tetramethyl-p-phenylenediamine dihydrochloride (TMPD)-ascorbate (0.9:9mM) to respiration solution. Rates of respiration are given in nmoles O2/min/mg dry weight. The respiratory acceptor control ratio (ACR) was calculated from basal mitochondrial respiration rate (in the presence of pyruvate/malate 1/4 mM, without ADP) and oxygen consumption after addition of 2mM ADP.

### Enzyme activity

Frozen tissue samples were weighed, homogenized (Bertin Precellys 24) in ice-cold buffer (50 mg/ml) containing 4-(2-hydroxyethyl)-1-piperazineethanesulfonic acid (HEPES) 5 mM (pH 8.7), EGTA 1 mM, DTT 1mM and 0.1% Triton X-100. Enzymatic activities were measured spectrophotometrically using standard coupled enzymes assays at 30°C. Citrate synthase (CS) activity was measured in a Tris-HCl buffer (pH 8) containing acetyl-Coa (10 mM), oxaloacetic acid (10 mM) and 5,5’-dittiobis-(2-Nitrobenzenoic acid) (DTNB) that allows, through coupled reactions involving CS, the production of the C_6_O_4_S_2_^-^ ion the apparition of which was followed at 412 nm. Complex I (NADH-CoQ reductase) activity was determined by monitoring the disappearance of NADH at 340 nm in a 50 mM phosphate buffer (KH_2_PO_4_ (0.411 mM), K_2_HPO4 (0.088 mM, pH 7.4) containing 3.75mg/ml BSA, 100 μM NADH and 100 μM of decyl-ubiquinone that played the role of electron acceptor. Complex IV (cytochrome oxidase complex) activity was assessed in a 90% reduced cytochrome c phosphate buffer (K_2_HPO4 50mM, cytochrome 1 mM, pH 7.4) by following reduced cytochrome c disappearance at 550 nm. Results are given in IU/g protein.

### Quantitative Real-time PCR

Frozen tissue samples were weighed and homogenized (Bertin Precellys 24) in Trizol reagent (Invitrogen) allowing total ventricular RNA extraction. After addition of chloroform, samples were centrifugated at 10.000 g at 4°C to obtain three phases: phenol-chloroform fraction, interphase and aqueous phase containing proteins, DNA and RNA respectively. Total RNAs were then precipitated by the addition of isopropanol to the aqueous phase, centrifugated at 10.000 g and resuspended in DNase/RNase free water. Oligo-dT first strand cDNA was synthesized from 1 μg of total RNAs using iScript cDNA synthesis kit (Bio-rad). Negative controls without reverse transcriptase enzyme were performed to verify the absence of genomic DNA in the samples. Real-time PCR was performed using the SYBR^®^ green method on CFX96 TouchTM Real Time PCR Detection system (Bio-Rad) from 2.5 ng cDNA. mRNA levels for all target genes were normalized to Tyrosine 3-Monooxygenase/Tryptophan 5-Monooxygenase Activation Protein Zeta (YWHAZ) expression level. Primer sequences are available in Table S1.

### Vitamin B12 dosage

Frozen myocardium tissue samples were weighed and homogenized in HCL (1N). Two-step immunoassay for the quantitative determination of vitamin B12 using chemiluminescence microparticle immunoassay (CMIA) on Alinity i ^®^ multiparametric instrument (Abbott, IL, USA) was performed.

### Liquid Chromatography-Mass Spectrometry

A liquid-chromatography coupled to mass spectrometry analysis was developed and carried out to quantified Methyl-NAM.

#### Tissue preparation

After sampling, heart has been reduced to a powder and 30 mg were weighed. The extraction was carried out using 800 μl of ice-cold 80% methanol. This is a semiquantitative method based on the use of deutered Nicotinamide (NAM-d4, N407752 Toronto Research Chemicals, Canada), added at the final concentration of 1mM. The tissue was homogenized in the Tissue Lyser (MBIZR-0209-99, MBI Lab Equipment, Canada) at 20 Hz (speed 5 during 45 seconds). After a centrifugation of 15min at 16,000 rpm (Sigma 4-16K, MBI Lab Equipment, Canada), the supernatant was collected and evaporated overnight in Speed Vac (RVC 2-33 CDplus, MBI Lab Equipment, Canada) at 5 mbar overnight. The sample was reconstituted with 100 μl of 60% of acetonitrile (A955-4, fisher chemical, Canada) diluted in water. After a sonication step (M1800, Emerson Branson, Canada) and a rapid centrifugation step of 2.5 min at 2000 rpm (Sigma 2-7, MBI Lab Equipment, Canada), the sample was transferred into a glass vial and placed in the HPLC coupled with a 6495 triple quadrupole MS/MS system (Agilent Technologies, USA) for the LC-MS analysis.

#### LC-MS analysis

Standards and samples were injected onto Agilent InfinityLab Poroshell 120 HILIC-Z column, 2.1 mm × 150 mm, 2.7 μm, PEEK-lined combined with UHPLC Guard Infinity Lab Poroshell HILIC-Z 2.1 x 5 mm, 2,7m (Agilent 821725-947) at the volume of 6 μl. For the high pH method, mobile phase A was consisted of ammonium acetate [10 mM, pH 9] adjusted with ammonium hydroxide. Mobile phase B was composed of 85% of acetonitrile combined with ammonium acetate [10 mM, pH 9] adjusted with ammonium hydroxide with a flow rate of 250 μl/min at a controlled temperature of 25°C. To minimize the binding of polar ionic metabolites to trace levels of metal in the system, InfinityLab Deactivator Additive (p/n 5191-4506, Agilent Technologies, USA) was added to the high-pH mobile phase. The analysis has been done in positive mode. The following linear gradient was run for 20 min (0–3min 98% B, at 11 min 30% B, 12min 40% B, 16-18 min 95%B, 18-20min 2% B). To determined retention times of Methyl-NAM, a standard molecule was used (1-Methylnicotinamide chloride (MeNAM) Sigma M4627). Retention times are shown in Table S2. Relative semi-quantitative values were determined using area ratios of the internal standard NAM-d4. Data were processed and analyzed with the Mass Hunter Qualitative Analysis (Agilent Technologies, version B.06.00). The reproducibility of the semi-quantification method was evaluated by calculating inter- and intra-assay coefficients of variability (CV) in 30 mg of control dog cardiac tissue powder (Table S3).

### Statistical analysis

The results are presented as the means ± standard error of the mean (SEM), with individual values for each condition. Statistical analysis was performed using GraphPad Prism 9 software (GraphPad Software, RITME, France). The significance of the differences between groups was evaluated first after verifying the normality of the distribution by a Shapiro-Wilk normality test. If the distribution was normal, an ordinary one-way analysis of variance (ANOVA) followed by Tukey’s multiple comparisons test was done. If it was not normal, a nonparametric Kruskal-Wallis test followed by an uncorrected Dunn’s test was chosen. To compare significance on two parameters such as treatment and sex, two-way analysis of variance (ANOVA) followed by uncorrected Fisher’s LSD test was done. P values ≤ 0.05 were considered statistically significant. For the survival curve, to compare two groups, a Logrank (Mantel-Cox test) analysis was done.

## Results

### 1. The 3VitB cocktail improves survival and physical capacities of mice subjected to cardiac pressure overload

Cardiac pressure overload induced mortality over time in mice fed with non-supplemented diet (ND) or diet supplemented with 3VitB cocktail when compared with sham groups in both sexes (Figure 1A). Survival was significantly improved in males fed with 3VitB as shown by Kaplan-Meier curve and the significant increase in median survival (Figure 1B). These effects were combined with a beneficial impact on the normal body weight gain in males with ageing that was impaired in male TAC-ND group (Figure 1C). In females, in spite of a trend towards a lower mortality rate and a higher median survival in TAC-3VitB group in comparison with TAC-ND group, it remained below study’s defined significance level (Figure 1A). This difference might be explained by the higher median survival in TAC-ND group in females than in males (Figure1B and supplementary table S3, p=0.03). Also of note, TAC did not affect weight gain in females (Figure 1C).

**Figure 1:**
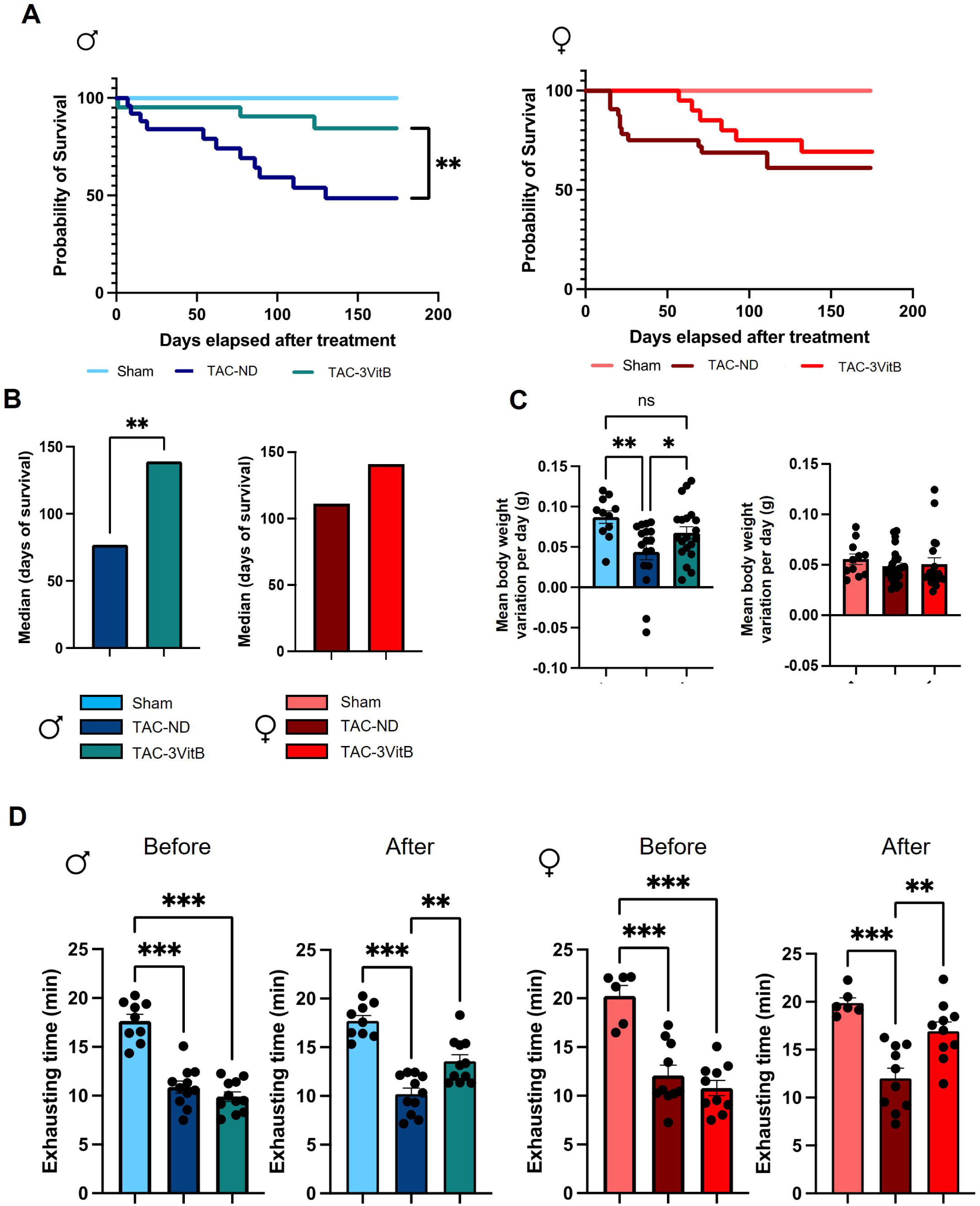
The 3VitB cocktail improves survival and physical capacities of mice subjected to cardiac pressure overload. **A**. Survival curve analysis of males and females treated with ND or 3VitB compared using log-rank test. **B.** Median survival after diet introduction. **C.** Body weight gain from surgery to death or to sacrifice 24 weeks after surgery**. D.** Maximal of running time on increasing speed on treadmill before and 8 weeks after treatment introduction. (n=11 to 20), ** p<0.01, *** p<0.001.

Exercise stress testing is an important noninvasive test to diagnose and risk stratify HF in patients. Although analysis of exercise capacity before treatment at 4 weeks after TAC showed an equal reduction in the animals randomly assigned to ND and 3VitB treatment, in females as well as in males, exercise capacity was significantly improved by 8 weeks of 3VitB treatment in both sexes and females showed higher capacities than males (Figure 1D, Table S3, p=0.003).

### 2. Cardiac function of mice affected by HF is protected by the 3VitB cocktail, especially in males

Since physical capacities are used by New York Heart Association to stratify the stage of HF, the improved exercise capacity on treadmill in TAC-3VitB groups could suggest a better preservation of cardiac function after 3VitB treatment introduction in this model. Importantly, we verified and confirmed that the TAC procedure triggered the same level of constriction and pressure gradient across the aortic cross in males and females before treatment as assessed by pulse wave Doppler analysis (Supplementary Figure S2A and B and Table S3, p>0.99), resulting in similar increase in LV mass and reduction in LVEF in both sexes at 4 weeks before treatment (Supplementary Figure S2C), ruling out any differences in early TAC response as a cause of subsequent evolution in cardiac function.

In male TAC-3VitB group, the degradation of left ventricular ejection fraction (LVEF), stroke volume (SV) and the increase of LV end-systolic and -diastolic diameters over the time course of the survival study, were slower than in TAC-ND group (Figure 2A-D), thereby confirming a benefit of these vitamins on the preservation of LV function and remodeling. In females, LVEF, SV and LVDs were similarly affected in TAC-ND group as compared to males (Figure 2A-C and Table S3, p>0.05 between sexes in TAC-ND groups) although LVDd was less increased (Figure 2D, Table S3, p=0.01 between males TAC-ND and females TAC-ND). The 3VitB treatment did not improved these parameters in females. Interestingly, the evaluation of the late diastolic transmitral flow velocity (E/A ratio) showed a similar increase 8 weeks after TAC in males and females (Figure 2E and Table S3, p=0.17 between sexes in TAC-ND groups) indicative of LV diastolic dysfunction and 3VitB treatment reduced this ratio to normal levels in both sexes.

**Figure 2:**
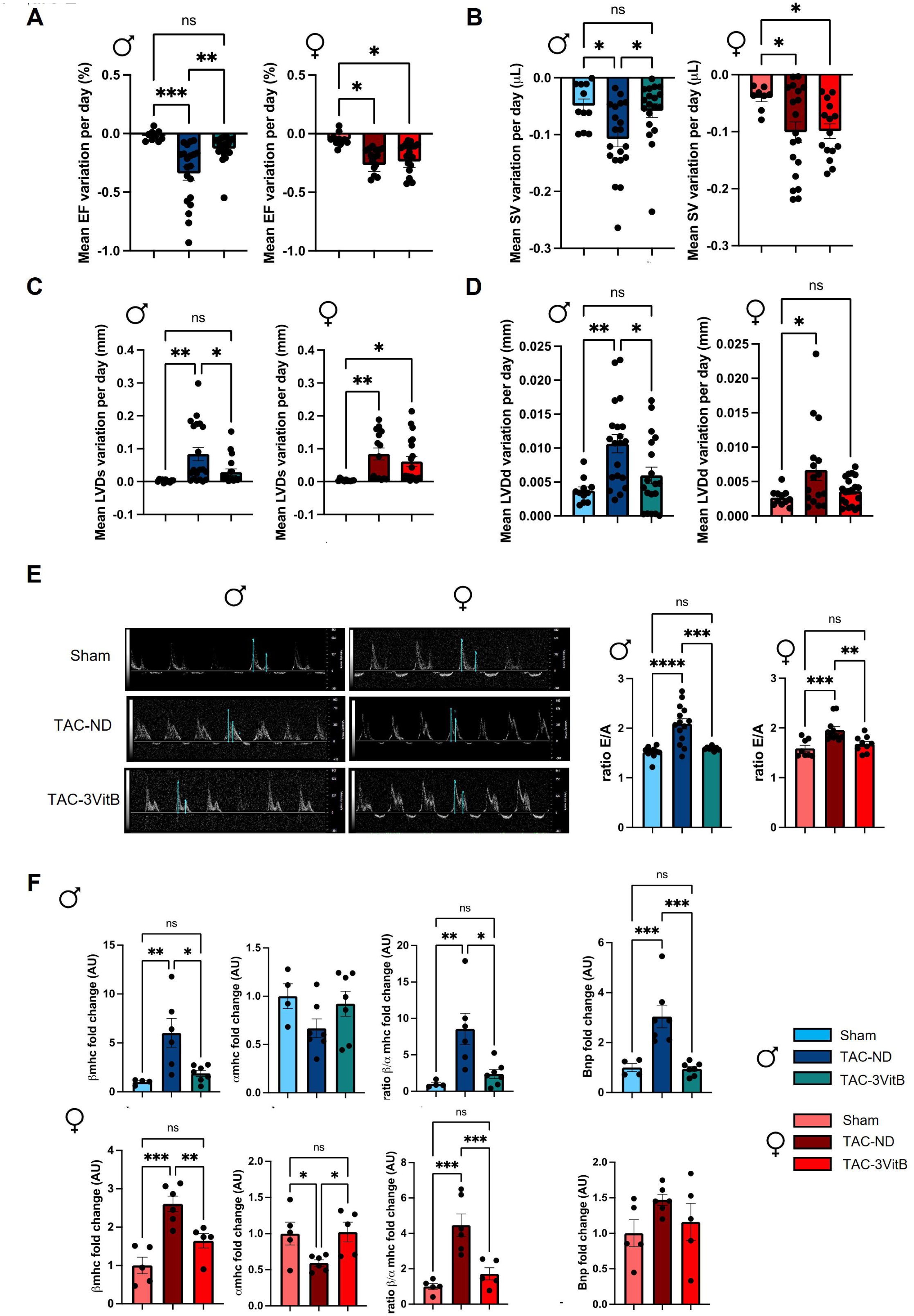
Cardiac protective effects of the 3VitB cocktail. **A.** Ejection fraction (EF) progression from surgery to death or to sacrifice 24 weeks after surgery. **B.** Stroke volume (SV) progression from surgery to death or to sacrifice 24 weeks after surgery **C.** End-systolic left ventricle diameter (LVDs) progression from surgery to death or to sacrifice 24 weeks after surgery. **D**. End-diastolic left ventricle diameter (LVDd) progression from surgery to death or to sacrifice 24 weeks after surgery **E.** Representative echocardiographs and E wave/A wave ratio (E/A ratio) 8 weeks after treatment introduction **F.** mRNA expression level of β-myosin heavy chain (*β-Mhc*), α-myosin heavy chain (*α-Mhc*) and brain natriuretic peptide (*Bnp*). (n=11 to 20), * p<0.05, ** p<0.01, *** p<0.001.

The quantification in LV myocardium of transcripts of molecular markers of HF in mice showed the characteristic shift from cardiac fast α-myosin heavy chain (*Mhc*) to slow *βMhc* and the increase in brain natriuretic peptide (*Bnp*) gene expression associated with LV dilation in males TAC-ND group; and the 3VitB treatment robustly reduced these markers (Figure 2F). In females, the *βMhc*/*αMhc* ratio increased less after TAC than in males (Figure 2F and Table S3, p=0.02 between sexes in TAC-ND groups) and it was rescued by the 3VitB treatment. *Bnp* expression was much less induced by TAC in females in agreement with the milder dilation of the LV chamber.

### 3. The 3VitB cocktail prevents the development of cardiac hypertrophy and fibrosis induced by cardiac pressure overload

In order to study the potential effects of the 3VitB cocktail on cardiac structure, a series of animals was especially designed to be sacrificed after 8 weeks of B vitamins supplementation, i.e. 12 weeks after TAC (Figure S1B). In males, heart weight-to-tibia length ratio (HWTL) at sacrifice was significantly increased in TAC-ND group and reduced in the TAC-3VitB group (Figure 3A). In females, HWTL was significantly increased in TAC-ND group but much less than in males and the 3VitB treatment did not further reduced this parameter (Figure 3A, and Table S3, p<0.001 p=0.02 between sexes in TAC-ND groups). In contrast, TAC increased lung weight-to-tibia length ratio in both sexes suggesting similar levels of pulmonary oedema and the 3VitB had same beneficial impact in both sexes (Figure 3B). In spite of the lower increase of HWTL ratio triggered by TAC in females compared to males, the mean cross-sectional area of cardiac fibers and the proportion of very thick cardiomyocytes showed similar and significant increase in both sexes in TAC-ND groups (Figure 3C and D). This confirmed the activation of the hypertrophy process in these groups that was counterbalanced by 3VitB treatment as judged by the “treatment effect” revealed by the 2-way ANOVA statistical analysis including males and females together (Table S3, treatment factor p <0.001). This histological analysis was completed with Sirius red staining that showed a significant rise in cardiac fibrosis in TAC-ND group in males that was prevented by the 3VitB supplementation (Figure 4A and B). This observation is reinforced by the measurement of the expression of collagen I that showed a significant increase in the TAC group fed with ND (Figure 4C) while no increase was reported when mice were under 3VitB treatment. No expression change in collagen III was measured in TAC groups. In females, no such increase in cardiac fibrosis and collagen I expression was observed in the female TAC-ND group (Figure 4A, B and C).

**Figure 3:**
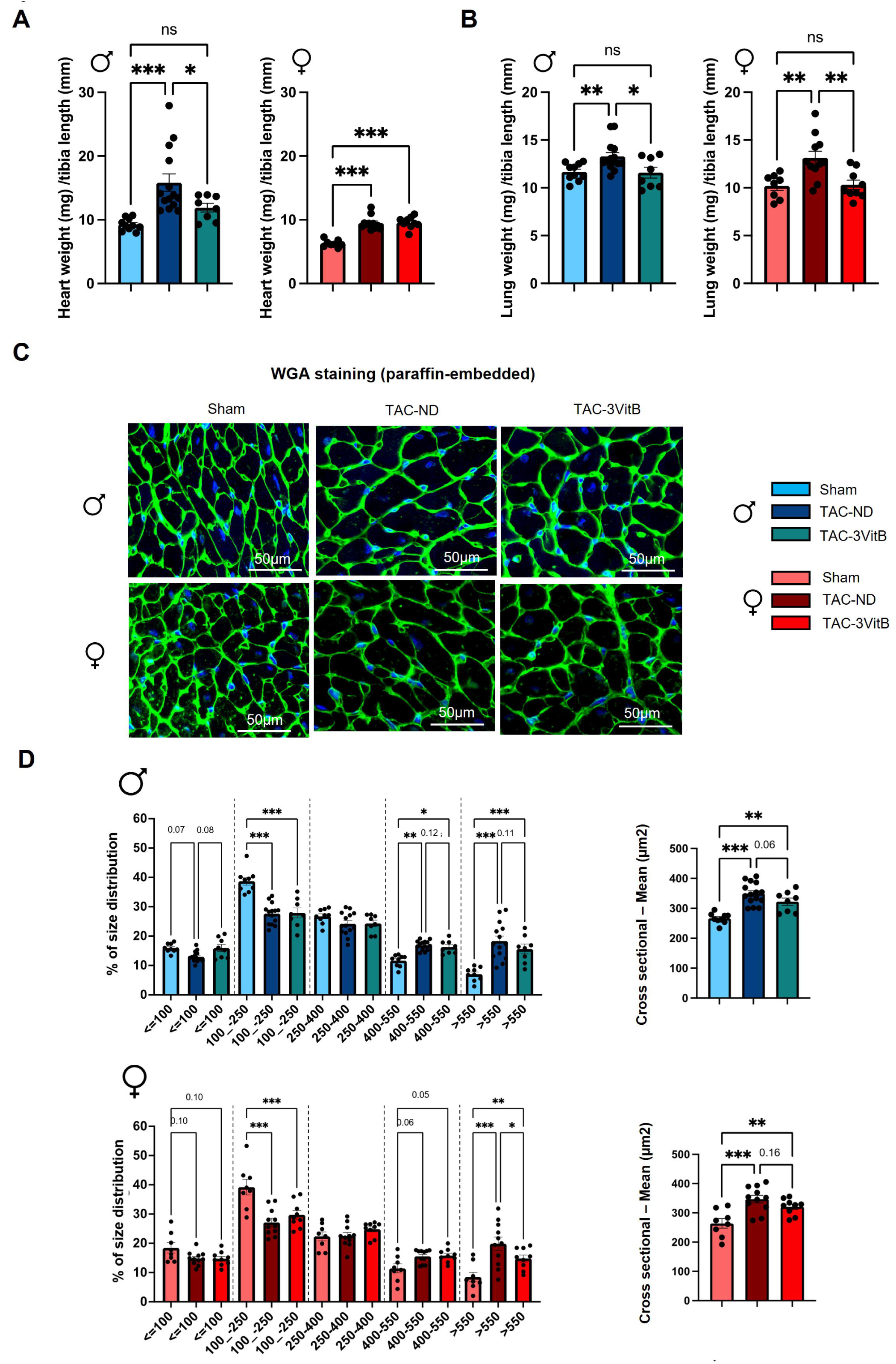
The 3VitB cocktail prevents the development of cardiac hypertrophy. **A.** Heart weight to tibia length ratio (HWTL) at sacrifice 8 weeks after 3VitB treatment introduction. **B.** Lung weight to tibia length ratio (LXTL) at sacrifice 8 weeks after 3VitB treatment introduction. **C.** Representative pictures of cardiomyocyte hypertrophy analysis by fluorescein isothiocyanate (FITC)-labeled wheat germ agglutinin (WGA) after 8 weeks of 3VitB treatment. **D.** proportion of cardiomyocytes by cross sectional size category, mean cross sectional area. (n=6 to 8 animals per group), * p<0.05, ** p<0.01, *** p<0.001.

**Figure 4:**
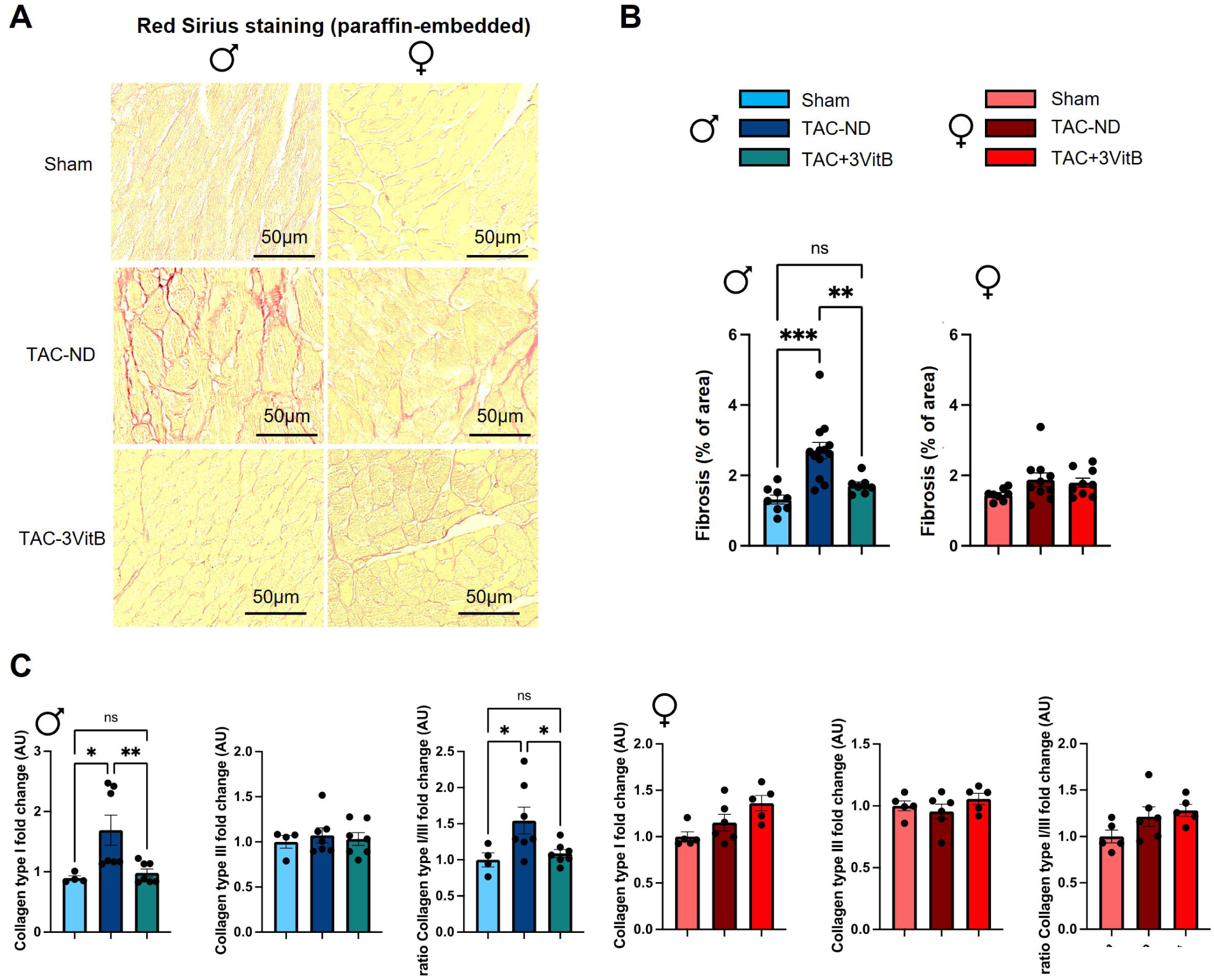
Myocardial fibrosis after 8 weeks of treatment. **A.** Representative pictures of myocardial fibrosis analysis by Sirius red after 8 weeks of 3VitB treatment. **B.** Proportion of total fibrosis in the different experimental groups at sacrifice 8 weeks after 3VitB treatment introduction. **C.** Left ventricular mRNA expression level of collagen I (*Col1a*) and collagen III (*Col3a*). (n=6 to 8 animals per group), * p<0.05, ** p<0.01, *** p<0.001.

### 4. Energy metabolism of mice affected by HF is preserved by the 3VitB cocktail, especially in males

Considering the major role of B vitamins in energy metabolism and mitochondrial metabolism in particular, key elements of energy metabolism were characterized in the myocardium of the animals sacrificed 8 weeks after the beginning of the treatment. In males, mitochondrial oxidative capacities measured in the presence of substrates feeding complex I and II and direct activation of complex IV were altered in the TAC-ND group and fully rescued by the 3VitB treatment (Figure 5A). In females, mitochondrial respiration rates were only mildly affected in the TAC-ND group, although they were not significantly different than in TAC-ND males (Table S3). Yet they were significantly increased by the 3VitB treatment.

**Figure 5:**
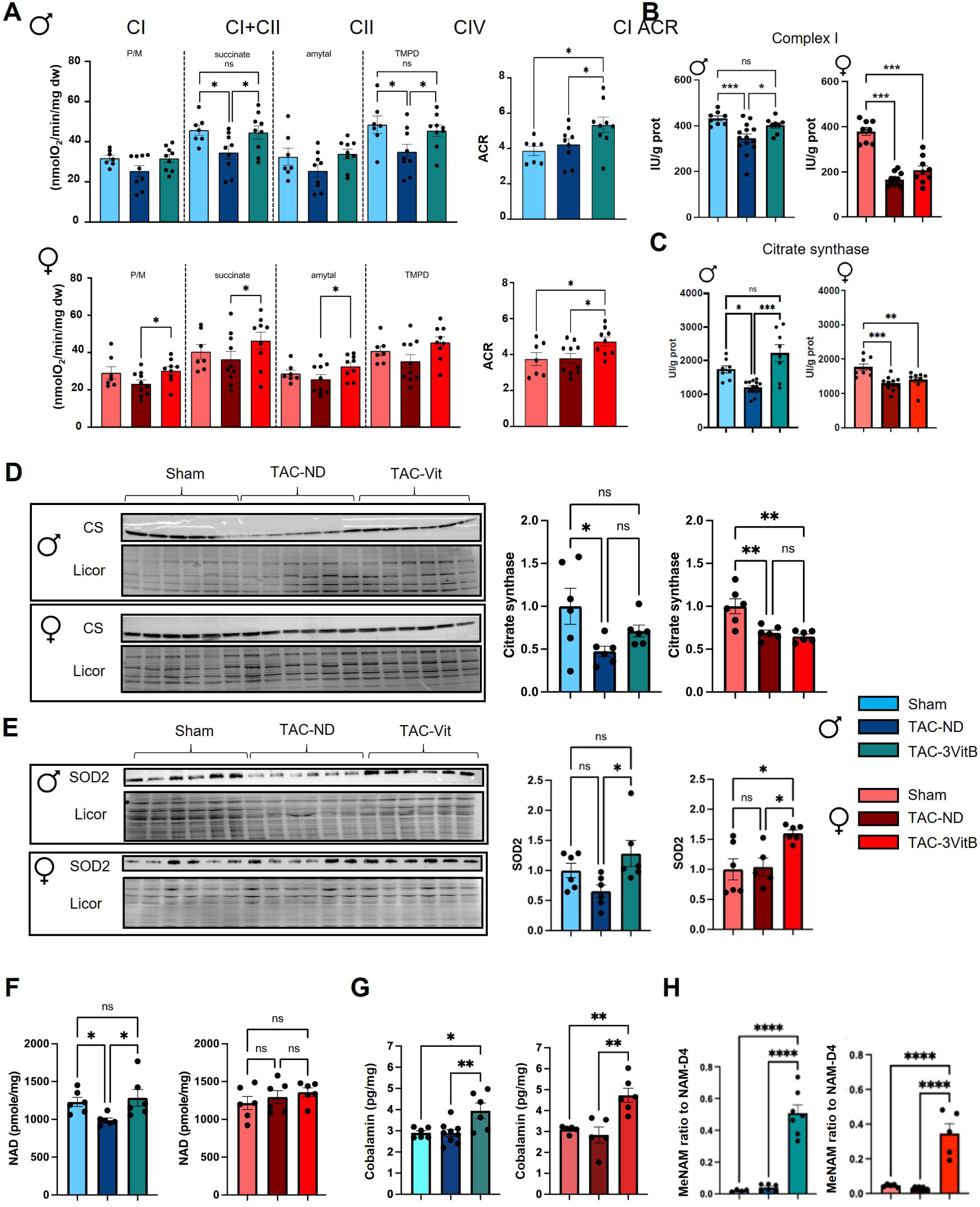
Cardiac mitochondrial oxidative capacities of mice affected by HF is preserved by 8 weeks of 3VitB cocktail treatment. **A.** Rate of respiration measured under phosphorylating conditions (2mM ADP) in the presence of substrates for complex I (P/M), complexes I+II (succinate), complex II (Amytal) and complex IV (TMPD). **B.** Complex I enzymatic activity. **C.** Citrate synthase (CS) enzymatic activity. **D.** Citrate synthase (CS) protein level. **E.** SOD2 protein level. **F.** Myocardial NAD level. **G.** Myocardial cobalamin level. **H.** Myocardial Methyl-NAM (MeNAM) level. (n=6 to 8 animals per group), * p<0.05, ** p<0.01, *** p<0.001.

Interestingly, the beneficial impact of the 3VitB treatment on mitochondrial oxidative capacities was also revealed by the higher respiratory acceptor control ratio (ACR) as measured in NAD-dependent complex I driven respiration in both males and females in comparison with Sham and TAC-ND groups (Figure 5A). The decrease in mitochondrial oxidative capacities in TAC-ND males was associated with significant reduction of complex I biochemical activity (Figure 5B) and citrate synthase (CS) activity and protein level (Figure 5C and D), CS being usually used as a marker of mitochondrial mass; these alterations were prevented by 3VitB supplementation. In females, despite their lesser sensitivity to TAC, the activities of these enzymes and CS protein level were affected in TAC-ND groups (Figure 5B, C and D) and not restored by 3VitB cocktail. The measurement of the mitochondrial superoxide dismutase 2 (SOD2) protein level revealed a rise in protein level in both TAC-3VitB groups that suggests that the 3VitB treatment could promote an antioxidant system (Figure 5E). Higher oxidative capacities in TAC-3VitB males were associated with prevention of NAD level drop induced by TAC (Figure 5F). Incidentally, the assessment of NAD and cobalamin level in myocardium showed significant increases that demonstrated the assimilation of NR and vitamin B12 by the 3VitB treated mice (Figure 5F and G). Methyl-NAM metabolite that represents the end-pathway product of both vitamin B3 elimination (nicotinamide) and vitamins B12/9 action on methylation, was induced by the 3VitB diet in both sexes (Figure 5H).

### 5. The 3VitB cocktail sustained mitochondrial biogenesis transcription cascade in pressure overloaded myocardium in males and females

Although alterations and preservation of oxidative capacities were observed in TAC-ND and TAC-3VitB males respectively, no significant modulation of *Pgc-1α*, *Pgc-1β* and *Nrf1* transcripts level was noticed (Figure 6A). The expression of these three genes involved in mitochondrial biogenesis was not altered in TAC-ND females neither when compared to sham group and only a slight increase in *Nrf1* expression was noticed in TAC-3VitB female group in comparison with Sham and TAC-ND groups. The 3VitB cocktail globally normalized of the expression of mitochondrial transcription factor A (*Tfam*), cytochrome c oxidase 4 (*Cox4*) and medium-chain acyl-CoA dehydrogenase (*Mcad*) in both sexes, with some minor differences (Figure 6A). At the protein level, PGC-1α protein level was not altered by surgery or treatment but a significant increase in NRF1 protein level was observed in TAC-3VitB groups in both males and females (Figure 6B).

**Figure 6:**
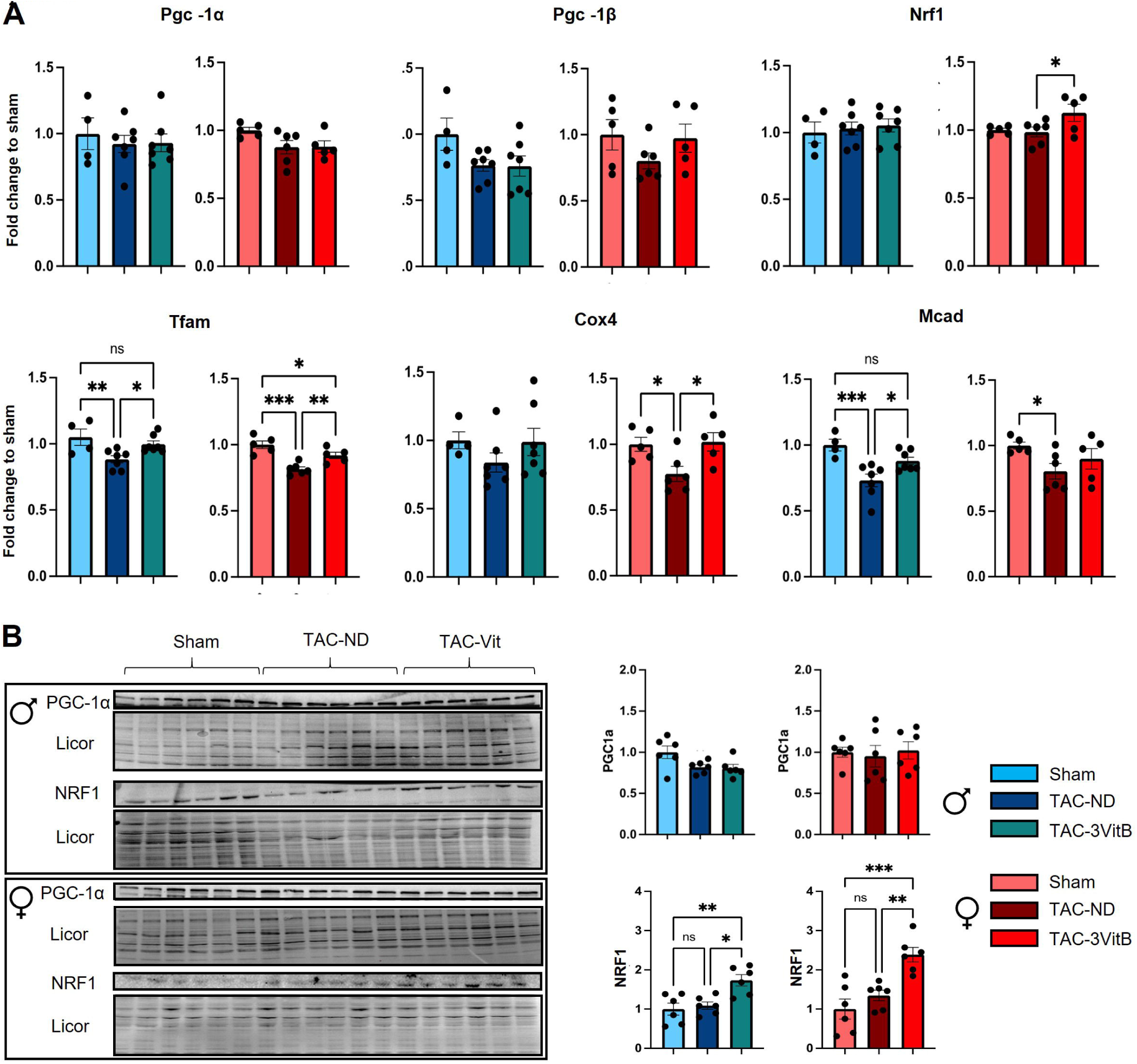
Cardiac mitochondrial oxidative capacities of mice affected by HF is preserved by 8 weeks of 3VitB cocktail treatment. **A.** mRNA expression level peroxisome proliferator-activated receptor gamma co-activator 1 α (*Pgc-1α*), peroxisome proliferator-activated receptor gamma co-activator 1 β (*Pgc-1β*), nuclear respiratory factor 1 (*Nrf1*), mitochondrial transcription factor A (*Tfam*), cytochrome oxidase subunit 4 (Cox4) and the medium-chain acyl-CoA dehydrogenase (*Mcad*). **B.** PGC-1α and NRF1 protein level.

While the increase in the expression of PGC-1α downstream cascade genes in male and female TAC-3VitB groups (Figure 6A) could explain the preservation of mitochondrial oxidative capacities (Figure 5A), the absence of any variation in PGC-1α gene expression and protein level between the different experimental groups suggested a post-translational activation of this transcriptional coactivator. Amongst the strongest activators of PGC-1α, SIRT1 requires NAD^+^ for its deacetylase activity which is thus largely dependent on NAD metabolism. 3VitB cocktail prevented the significant drop of NAD level induced by TAC in males while NAD level was not altered by TAC in females (Figure 5E). Whereas the vitamin treatment could help maintain NAD level, a prerequisite for optimal SIRT1 activity, this deacetylase did not show any significant changes in protein level in the different groups in both sexes, even though a trend towards a higher SIRT1 content was noticed in females after treatment (Figure 7A). The acetylation level of its target P53 tended to increase in both TAC-ND groups and was significantly reduced after 8 weeks of vitamins treatment in females only, revealing the activation of SIRT1 by the 3VitB cocktail in females (Figure 7A).

**Figure 7:**
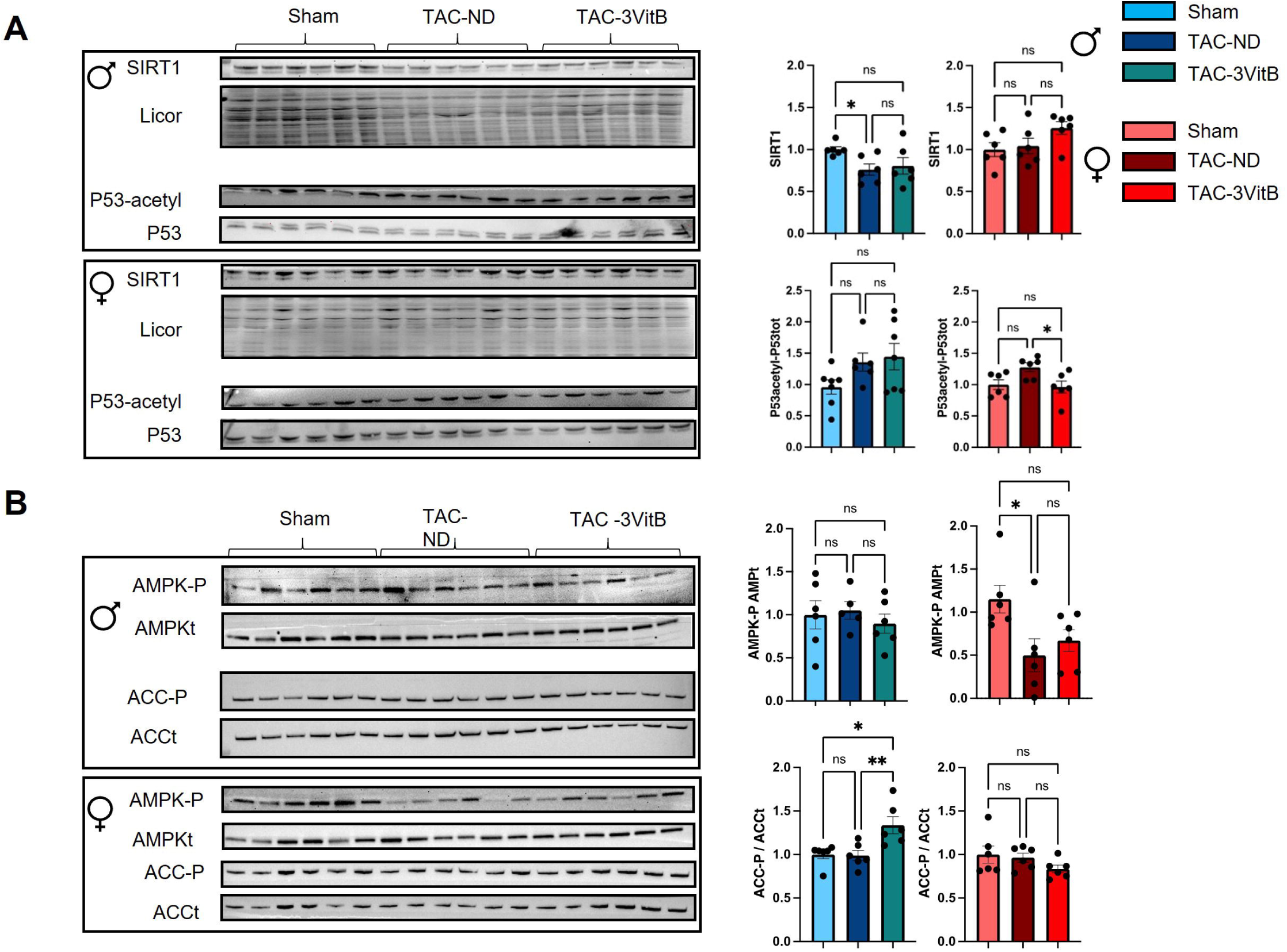
Cardiac mitochondrial oxidative capacities of mice affected by HF is preserved by 8 weeks of 3VitB cocktail treatment. **A.** Immunoblotting of PGC-1α, total-P53 (P53) and acetylated P53 (P53-acetyl). **B.** Immunoblotting of total AMPK (tAMPK), phosphorylated-AMPK (AMPK-P), total ACC (tACC)/phosphorylated-ACC (ACC-P).

As PGC-1α can be activated by AMP-activated protein kinase (AMPK) as well, a potential activation of this kinase was assessed by measuring phosphorylated-AMPK (P-AMPK) and phosphorylated-acetyl-CoA carboxylase (P-ACC), one of the AMPK targets traditionally used to reveal AMPK activation. Although, P-AMPK/total-AMPK ratio was similar in all male groups, TAC-3VitB group exhibited significantly higher P-ACC/total-ACC ratio after 8 weeks of 3VitB treatment, unveiling the activation of AMPK in this group (Figure 7B). In contrast, no such activation was noticed in females for which P-AMPK/total-AMPK ratio seemed to be negatively affected in both TAC groups and no variation of P-ACC/total-ACC ratio was measured between the three female groups (Figure 7B).

## Discussion

These last years, intensive research led in the field of HF has allowed a better understanding of the physiopathology of this syndrome. Although grey areas remain, the research community agrees that energy metabolism alterations in the myocardium plays important roles in cardiac decompensation and should be a target of future therapies (Bertero & Maack, 2018; Ventura-Clapier *et al*., 2011). Despite this, so far no metabolic therapy of HF is routinely used in clinics to specifically improve failing myocardium energy metabolism, especially mitochondrial functions (Schwemmlein *et al*, 2022). Based on the protective action of three B vitamins (NR, folate and cobalamin) on energy metabolism of the heart in animals facing cardiac pressure overload when administrated separately and preventively in our previous studies (Diguet *et al*., 2017; Piquereau *et al*., 2016), we tested the potential positive effects of a cocktail of these vitamins (3VitB) in a murine model of established HF to investigate its curative use. Here, we showed that the introduction of a diet supplemented with NR, B9 and B12 in mice with established cardiac dysfunction exhibits sex-dependent beneficial effects with stronger impacts in males although numerous beneficial effects were also seen in females. Precisely, this study shows that this combination of B3, B9 and B12 vitamins slows down the degradation of cardiac function induced by pressure overload, improves the physical capacities of the animals and protects myocardium energy metabolism, especially mitochondrial oxidative capacities; these effects are more pronounced in males with significant improvement of survival in these mice that are more severely affected by TAC in comparison with females.

In recent years, the studies aiming at understanding the metabolic alterations encountered in HF demonstrated that structural and functional alterations in mitochondria are associated with major deregulations of NAD homeostasis (Mericskay, 2016; Tannous *et al*, 2021). Beyond the fact that NAD as a coenzyme is required to support fuel substrate oxidation, TCA cycle and oxidative phosphorylation in mitochondria to generate ATP, loss of NAD homeostasis has also been reported to impact enzymes with NAD^+^-dependent activity, such as Sirtuins, that could be involved in cardiac disease progression (Horton *et al*, 2016; Sanz *et al*, 2019; Walker *et al*, 2021). It was therefore with the idea of proposing a treatment that could both stimulate the fundamental processes of energy metabolism and support cellular levels of NAD that the combination of B-vitamins tested in this study was developed, hypothesizing that the addition of the respective effects of these vitamins on mitochondrial biogenesis and NAD metabolism (Diguet *et al*., 2017; Piquereau *et al*., 2016) could constitute a new tool in the therapeutic arsenal for treating HF. While the significant reduction in mortality in males is a strong argument in favor of the efficacy of this B-vitamin cocktail in slowing the progression of the disease, the improvement in physical capacities observed in both sexes is also highly significant, since it demonstrates better adaptation of the treated animals when the body’s energy demand increases. By extrapolation, this would be akin to a better exercise test score in HF patients, a predominant indicator in determining the stage of HF (Lim *et al*, 2018). Moreover, as exercise training and cardiac rehabilitation are considered as major non-pharmaceutical beneficial approaches in the treatment of HF (Eleyan *et al*, 2025), the increased physical capacity conferred by the 3VitB cocktail could enhance the benefit of these approaches. Thus, the subtler effects of this treatment on mortality and cardiac systolic function in the females, resulting from a milder phenotype that was expected due to widely-reported cardioprotective mechanisms in this sex (Blenck *et al*, 2016), should not obscure its benefits in female mice. Incidentally, the improvement in diastolic function, the reduction in pathological β−MHC isoform expression, and the milder cardiomyocytes hypertrophy in TAC females after treatment, comparable to what was observed in males, demonstrates the sensitivity of the females to these vitamins. Interestingly, 3VitB treatment improved physical capacities in exercise stress test in females despite a lack of clear improvement of systolic cardiac function although the latter was less affected than males; however, we note that diastolic dysfunction as assessed by E/A was improved as in males. Obviously, physical exercise capacity integrates a number of regulatory physiological processes and we measured the cardiac function parameters at rest since limitation of the murine model precludes to perform echocardiography while exercising. Therefore, we cannot know if the 3VitB treatment improves cardiac adaptation to exercise or other parameters as well including vascular resistance as reported for NR (Cao *et al*, 2022), skeletal muscle bioenergetics or respiratory function, as suggested by the reduced level of pulmonary oedema obtained with the 3VitB treatment.

In the cardiac left ventricle, the study of the mitochondrial respiratory chain on permeabilized ventricular fibers clearly highlighted an improvement in the energy-producing capacity of mitochondria in treated males and females. In the latter, these improved oxidative capacities are not associated with any significant effects of the vitamins on citrate synthase enzyme activity, a marker of functional mitochondrial mass, that remains degraded in 3VitB treated females compared to sham females. This was not the case in males, where the 3VitB cocktail normalized the activity of this enzyme, suggesting again that the treatment acts through different pathways according to sex. However, the activation of the expression of genes that are under the control of PGC-1α such as *Tfam*, *Cox4* and *Mcad* (Garnier *et al*, 2003), shows that the support of mitochondrial oxidative capacities, involves mechanisms that stimulate genes involved in energy metabolism in both sexes. It is now well known that PGC-1α plays a central role in the regulation of energy metabolism and that its expression and activity can be modulated through many ways (Miller *et al*, 2019). The various mechanisms leading to the activation of this major regulator of energy metabolism and mitochondrial life cycle in particular could therefore be differently stimulated by the B vitamins in males and females in the present model. These mechanisms could involve SIRT1, which appears to be activated only in treated females. SIRT1 is a deacetylase whose actions on PGC-1α activity and on metabolism in general has been highlighted in several studies (Barcena *et al*, 2023; Li, 2013; Matsushima & Sadoshima, 2015), and it could therefore be at the heart of mechanisms protecting energy metabolism. Deregulation of pathways involving SIRT1 has been described in the pathophysiology of HF (Liu *et al*, 2023), and the effects of a combination of cobalamin and folate (Garcia *et al*, 2011; Piquereau *et al*., 2016) and NR (Hu *et al*, 2022) on the activity of this enzyme could play a crucial role in the benefits observed in females. Activation of the SIRT1 deacetylase is particularly conducive to the maintenance of mitochondrial function via its involvement in mitochondrial biogenesis mechanisms (Jeninga *et al*., 2010), and it has recently been shown that activation of a SIRT1-PGC-1α-PPARα pathway by NR promotes mfn2-dependent mitochondrial fusion leading to improved cardiac mitochondrial pool quality (Hu *et al*., 2022). In the present study, differences in the activation of this enzyme in males and females raise questions, especially as this does not appear to be correlated with cellular NAD levels, as may be the case in other studies (Hong *et al*, 2018; Zheng *et al*, 2019). While stabilization of SIRT1 by MeNAM could explain its higher activity (Hong *et al*, 2015), the lack of effect of a similar increase in MeNAM level in 3VitB males, compared to 3VitB females, suggests more complex regulatory mechanisms that could involve the hormonal status of the animals or sex-linked chromosomal effects (Ventura-Clapier *et al*, 2017). In males, the TAC induced-cardiac pressure overload leads to a marked decrease in myocardial NAD levels, which are restored by the vitamin cocktail, in line with the effects of NR described in the literature (Abdellatif *et al*, 2021; Diguet *et al*., 2017; Tong *et al*, 2021). While this increase in cellular NAD levels in treated males has no effect on SIRT1 activity, which does not appear to be impaired in TAC males, it could support the activity of other NAD-dependent enzymes. This could notably be the case for SIRT3, which is known to be a major regulator of mitochondrial activity via deacetylation of a number of mitochondrial proteins (Wang *et al*, 2023). Indeed, it was shown that SIRT3 could be at the heart of NR’s cardioprotective effects in stressed heart through its involvement in the inhibition of mitochondrial oxidative stress (Zhao *et al*, 2024). Activation of SIRT3 could also allow deacetylation of mitochondrial proteins such as OPA1 whose hyperacetylation has been demonstrated in the heart facing pressure overload and its deacetylation by SIRT3 can be cardioprotective (Samant *et al*, 2014). However, a SIRT3-dependent effect of NR has been challenged by a recent study led by Tian’s team on a model of established HF (Walker *et al*, 2023). This study demonstrates that NR is protective independently of mitochondrial protein deacetylation by SIRT3 and would instead involve stimulation of NADH redox-sensitive short-chain dehydrogenase/reductase proteins, mitochondrial ribonuclease P protein 2 in particular (Walker *et al*., 2023). In this work, the beneficial effects of NR were preserved in SIRT3KO mice exhibiting hyperacetylation of cardiac mitochondrial proteins under stress, supporting the conclusion of a study published earlier asserting that high acetylation level of mitochondrial proteome does not promote HF (Davidson *et al*, 2020). Although this could not be verified in our study, it is likely that the mechanism described by Tian’s team is involved in the effects of the 3VitB cocktail, given the commonalities in the experimental protocols. However, the remarkable reduction in mortality in 3VitB males was not observed in mice treated with NR alone (Walker *et al*., 2023), suggesting a net benefit from the addition of cobalamin and folate in our study. The protection of cardiac function conferred by these two vitamins, already suggested in previous studies (Hagar, 2002; Miao *et al*, 2024; Octavia *et al*, 2017; Piquereau *et al*., 2016; Qipshidze *et al*, 2010), would allow a synergistic action with NR via the activation of other pathways that would reinforce the consequences of the treatment, in particular on energy metabolism. Among these pathways, the stimulation of AMPK revealed in males in the present study, potentially by the combination of cobalamin and folate as it has already been demonstrated previously (Piquereau *et al*., 2016), could play a role in improving the survival of the animals included in this study. AMPK, a key metabolic sensor, is known for its cardioprotective effects (Russell *et al*, 2004; Sasaki *et al*, 2009; Zarrinpashneh *et al*, 2006) and may control the regulatory mechanisms of energy metabolism in a sex-specific manner (Grimbert *et al*, 2021). Its activation only in males could thus partly explain the differences observed between both sexes, its involvement in the sex-specific effects of the 3VitB cocktail will thus be the subject of a future work.

In summary, the present work shows, in line with our previous studies, that the oral supplementation with a cocktail mixing folate, cobalamin and NR improves heart function, mitochondrial function and survival in mice facing cardiac pressure overload with a higher sensitivity in males. These vitamins could be an initial tool for targeting metabolic alterations in HF and may be a new option to treat this disease. Despite the considerable work carried out during this study, the precise mechanisms involved in the benefits conferred by this treatment have not been clearly identified, and this remains a major limitation of our study. However, AMPK in males and SIRT1 in females could play key roles that will be investigated in future works. Of course, therapeutic use of this compounds will also require further studies to determine pharmacokinetics of these vitamins and to assess safety and efficacity in human before translation to clinics.

## Acknowledgments

We thank Valérie Domergue and the Animex team (animal experimentation), facility of the “Ingénierie et Plateforme au Sercice de l’Innovation Thérapeutique” UMS IPSIT of Université Paris Saclay. We also acknowledge Dr Renée Ventura-Clapier and Pr. Vladimir Veksler for continuous support.

## Author contributions

J.P. and M.M. designed the study. S.E.B., M.D., A.K., M.G., I.M., B.B., F.M-N., A.I and J.P. carried out the experimental design and execution of experiments. S.E.B., M.D., A.K., M.G., I.M., B.B., F.M-N., A.I, M.R. and J.P. analyzed the data. S.E.B., J.P., M.M., A.G., C.L. and M.R. contributed to data interpretation. J.P., M.M. and S.E.B. wrote the manuscript. J.P., M.M. and M.R. critically revised the manuscript.

