## Supplemental figures and supplemental tables for "A cocktail of B vitamins with nicotinamide riboside, folate and cobalamin preserves cardiac function and mitochondrial oxidative capacities in a mouse model of heart failure"

**Supplemental figure legends**

**Figure S1: Schematic experimental design of the study divided into two main batchs.**

**A.** Evaluation of 3VitB cocktail impact on survival, cardiac function, and effort tolerance at 20 weeks of treatment. The randomization was done at 4 weeks after TAC surgery based on three inclusion criteria (pression gradient, LVEF, and LV mass). Mice male and female were divided into three groups: 1) sham-operated mice receiving standard non-synthetic diet (n=11 male and n=11 female Sham), 2) TAC mice receiving standard non-synthetic diet (n=20 male and n=16 female TAC-ND), 3) TAC mice receiving supplemented diet with NR, cobalamin and folate (n=19 male and n=20 female TAC-3VitB). Cardiac function was monitored by echocardiography at four weeks intervals.

**B.** Evaluation of 3VitB cocktail impact on cardiac tissue, HF and fibrosis biomarkers, mitochondrial function and NAD metabolomic evaluation at 8 weeks of treatment. The randomization was done at 4 weeks after TAC surgery. Mice male and female were divided into three groups: 1) sham-operated mice receiving standard non-synthetic diet (n=9 male and n=6 female Sham), 2) TAC mice receiving standard non-synthetic diet (n=11 male and n=10 female TAC-ND), 3) TAC mice receiving supplemented diet with NR, cobalamin and folate (n=11 male and n=10 female TAC-3VitB).

**C.** Throughout the study, especially for survival monitoring, an assessment of state of health using this scoring grid was carried out. The animal reaching a sum of the scores superior or equal to 5, or a score of 3 for only one of the 3 criteria was euthanized.

**Figure S2: Validation of TAC induced-pressure overload model at four weeks after surgery and mice randomization by echocardiography**

**A.** Four weeks after the surgery, the TAC model was confirmed for sham and TAC male and female mice by measuring the aorta diameter by echocardiography trans-thoracic (Vevo 3100, FUJIFILM Visualsonics Inc., Toronto, Canada) in Bidimensional mode (B-mode). The dilation of the aorta before the ligature (site A) and the reduction of the aorta after the ligature (site B) was measured. The view of the aortic cross structure was confirmed by an observation of the aortic flow in color doppler.

**B.** Around the ligature site, the aorta flow was disturbed and considerably accelerated in TAC model. Pressure gradient was measured by pulsed wave (PW) Doppler for sham and TAC male and female mice using the modified Bernoulli equation (Pressure gradient = 4*Velocity2). Images were analyzed by VevoLab Software (FUJIFILM Visualsonics Inc., Toronto, Canada). Mice with a pressure gradient higher than or equal to 60mm Hg were included in the study.

**C.** TAC induced-pressure overload model was correlated with an alteration of cardiac function and a cardiac hypertrophy. Ejection Fraction (LVEF) and left-ventricle mass (LV mass) were measured by echocardiography in parasternal long and short axis B-mode and M-mode to refine mouse randomization. TAC mice with a reduction of 10% of LVEF and an improvement of 30% were included in the study.

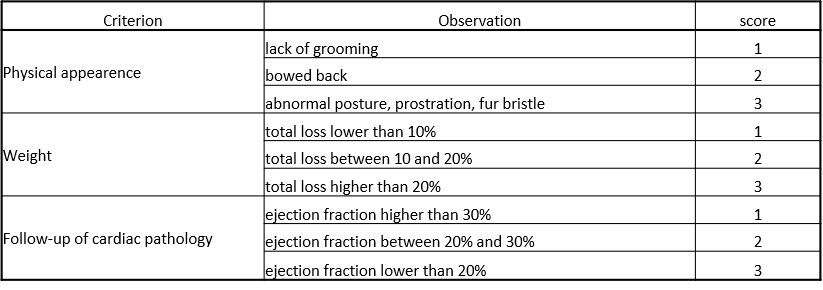

Echo at

**3W post TAC**

TAC

**Inclusion criteria**

Pression gradient

LVEF -10%

LVmass > 30%

4 weeks

Echo

**8 weeks**

Echo

**baseline**

Echo

**4 weeks**

**TAC+3VitB**

**Untreated SHAM**

**C57Bl6N**

n=11

n=20

n=19

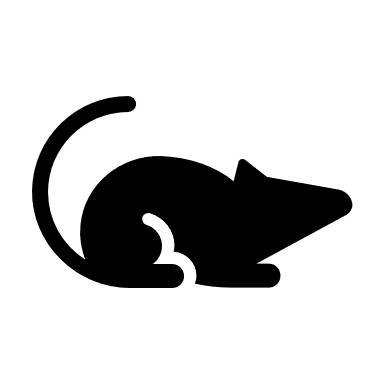

**C57Bl6N**

n=11

n=16

n=20

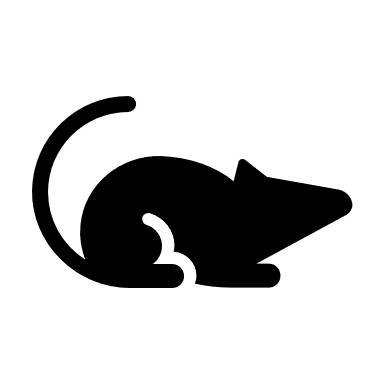

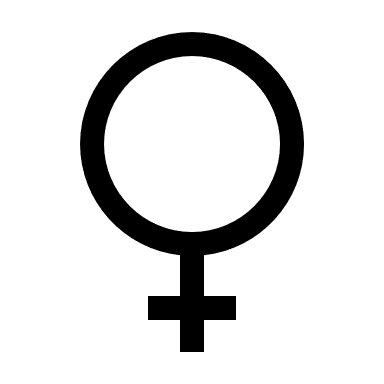

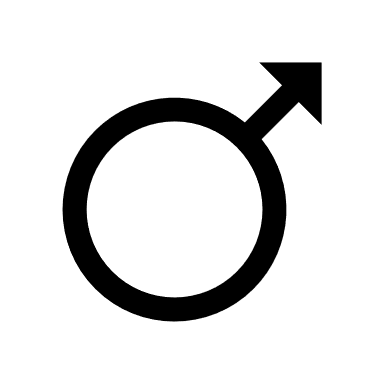

Randomization at 4W post TAC

TREATMENT

Echo

**20 weeks**

**TAC+ND**

**TAC+3VitB**

**Untreated SHAM**

**TAC+ND**

**Timeline BATCH 1**

**Inclusion criteria**

Pression gradient

LVEF -10%

LVmass > 30%

**TAC+3VitB**

**Untreated SHAM**

**C57Bl6N**

n=9

n=11

n=11

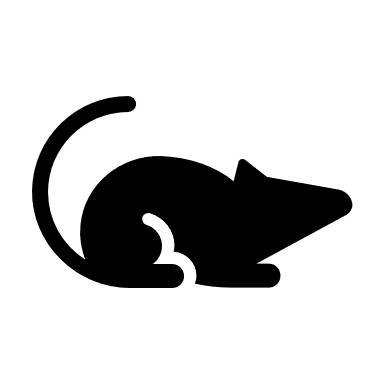

**C57Bl6N**

n=6

n=10

n=10

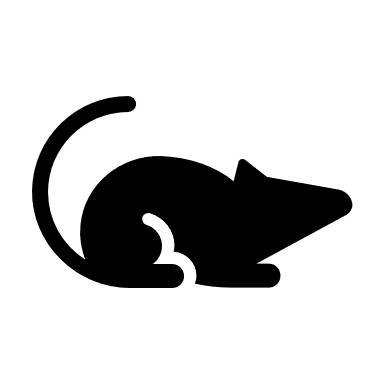

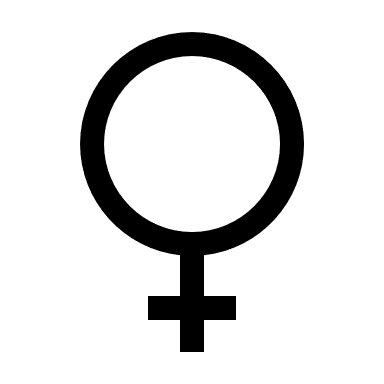

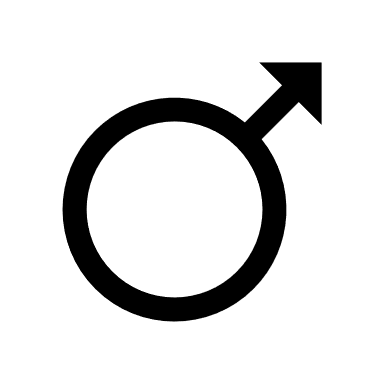

**TAC+ND**

**TAC+3VitB**

**Untreated SHAM**

**TAC+ND**

Echo

**16 weeks**

- Survival study stopped at 20 weeks of treatment
- Cardiac function monitoring
- Effort tolerance test before and 8 weeks after treatment

Randomization at 4W post TAC

Echo at

**3W post TAC**

TAC

4 weeks

Echo

**baseline**

TREATMENT

- Effort tolerance test before and 8 weeks after treatment
- Euthanized and tissus sampling at 8 weeks of treatment
- Cardiac function monitoring
- Relative quantification of heart failure and fibrosis biomarkers
- Histological analysis
- Mitochondrial function study
- NAD metabolomic evaluation

A

**Timeline BATCH 2**

B

Echo

**12 weeks**

Echo

**8 weeks**

Echo

**4 weeks**

C

Figure S1

After evaluation of state of health using this scoring grid, the animal reaching a sum of the scores superior or equal to 5, or a score of 3 for only one of the 3 criteria was euthanized.

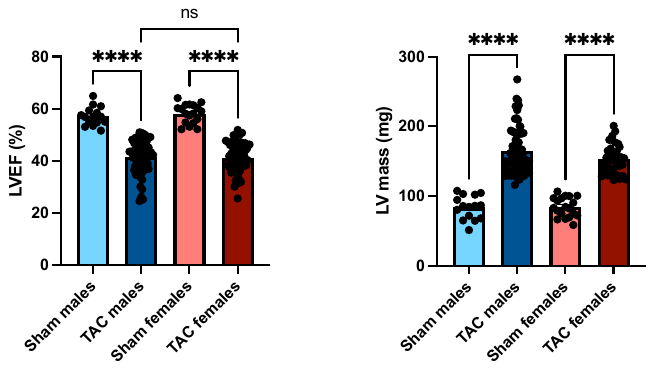

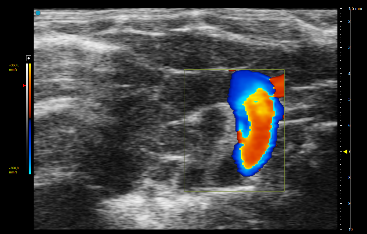

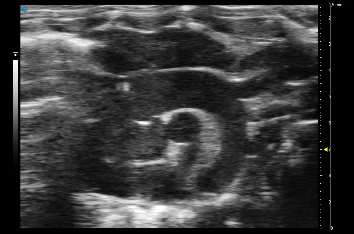

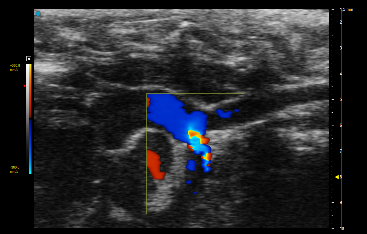

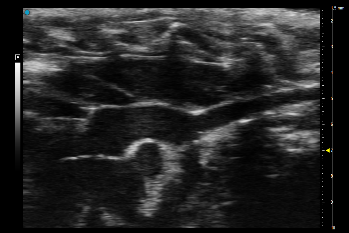

B-Mode

Color Doppler

Sham

TAC

**Male**

**Female**

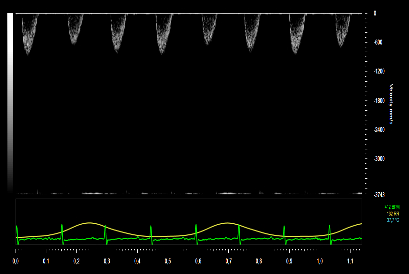

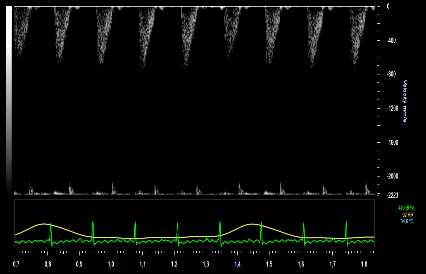

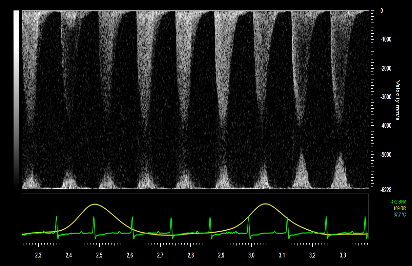

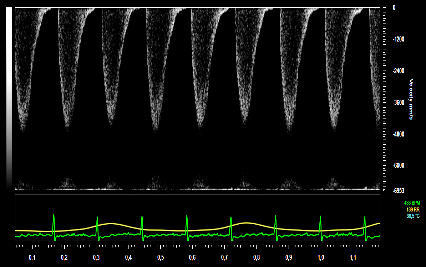

Sham

TAC

A

B

A

B

Pression gradient evaluation

Aortic diameter

Cardiac function

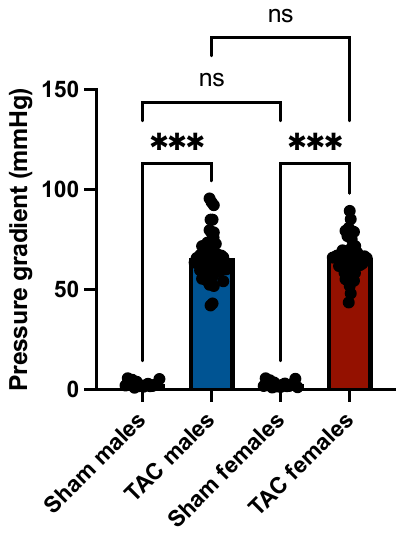

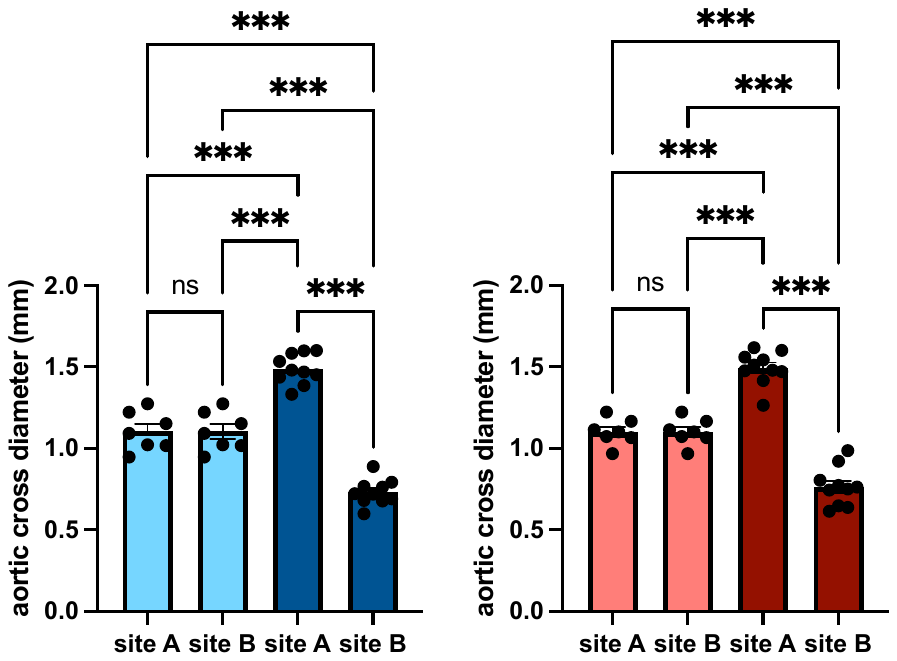

**Sham**

**Sham**

**TAC**

**TAC**

**A**

**B**

**C**

Figure S2

**Supplemental tables**

**Table S1.**

| **Gene** | **Primers** | **Hybridation temperature** |
| --- | --- | --- |
| *Col1a* | 5'-CTCAAGATGTGCCACTCTGACT-3' | 60°C |
|  | 5'-CTCCATGTTGCAGTAGACCTTG-3' |  |
| *Col3a* | 5'-GAT GGAAACCCTGGATCAGA -3' | 60°C |
|  | 5'-GCACCAGGAGAACCATTTTC-3' |  |
| *Cox4* | 5’-TGG GAG TGT TGT GAA GAG TGA -3’ | 57°C |
|  | 5’-GCA GTG AAG CCG ATG AAG AAC-3’ |  |
| *α-Mhc* | 5'-CCAATGAGTACCGCGTGAA-3' | 58°C |
|  | 5'-ACAGTCATGCCGGGATGAT-3' |  |
| *β-Mhc* | 5'-ATGTGCCGGACCTTGGAA-3' | 60°C |
|  | 5'-CCTCGGGTTAGCTGAGAGATCA-3' |  |
| *Mcad* | 5'-CCGTTCCCTCTCATCAAAAG-3' | 60°C |
|  | 5'-ACACCCATACGCCAACTCTT-3' |  |
| *Nrf2* | 5'-CACTCAACATTTCGGGAAGAG-3' | 60°C |
|  | 5'-CTCATTCATCTGTTGCTCTTGG-3' |  |
| *Pgc-1α* | 5'-CACCAAACCCACAGAGAACAG-3' | 58°C |
|  | 5'-GCAGTTCCAGAGAGTTCCACA-3' |  |
| *Pgc-1β* | 5'-TGGAAAGCCCCTGTGAGAGT-3' | 60°C |
|  | 5'-TTGTATGGAGGTGTGGTGGG-3' |  |
| *Tfam* | 5'-GCTAAACACCCAGATGCAAA-3' | 60°C |
|  | 5'-TACTTGCTCACAGCTTCTTTGT-3' |  |

Primers used for quantification of mRNA expression level.

**Table S2**

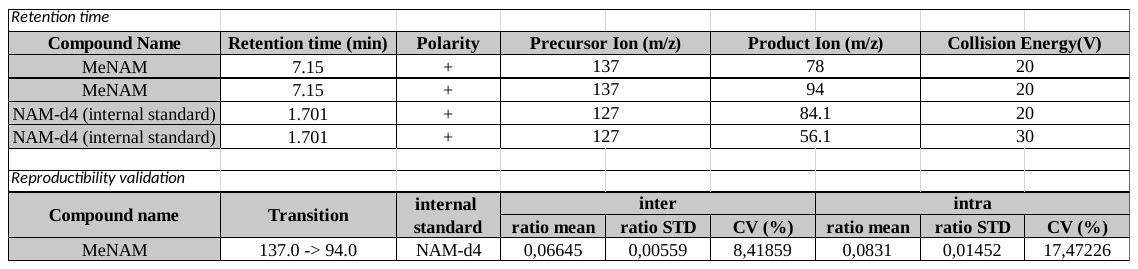

LC-MSMS. Retention time and Method reproducibility validation

**Table S3**

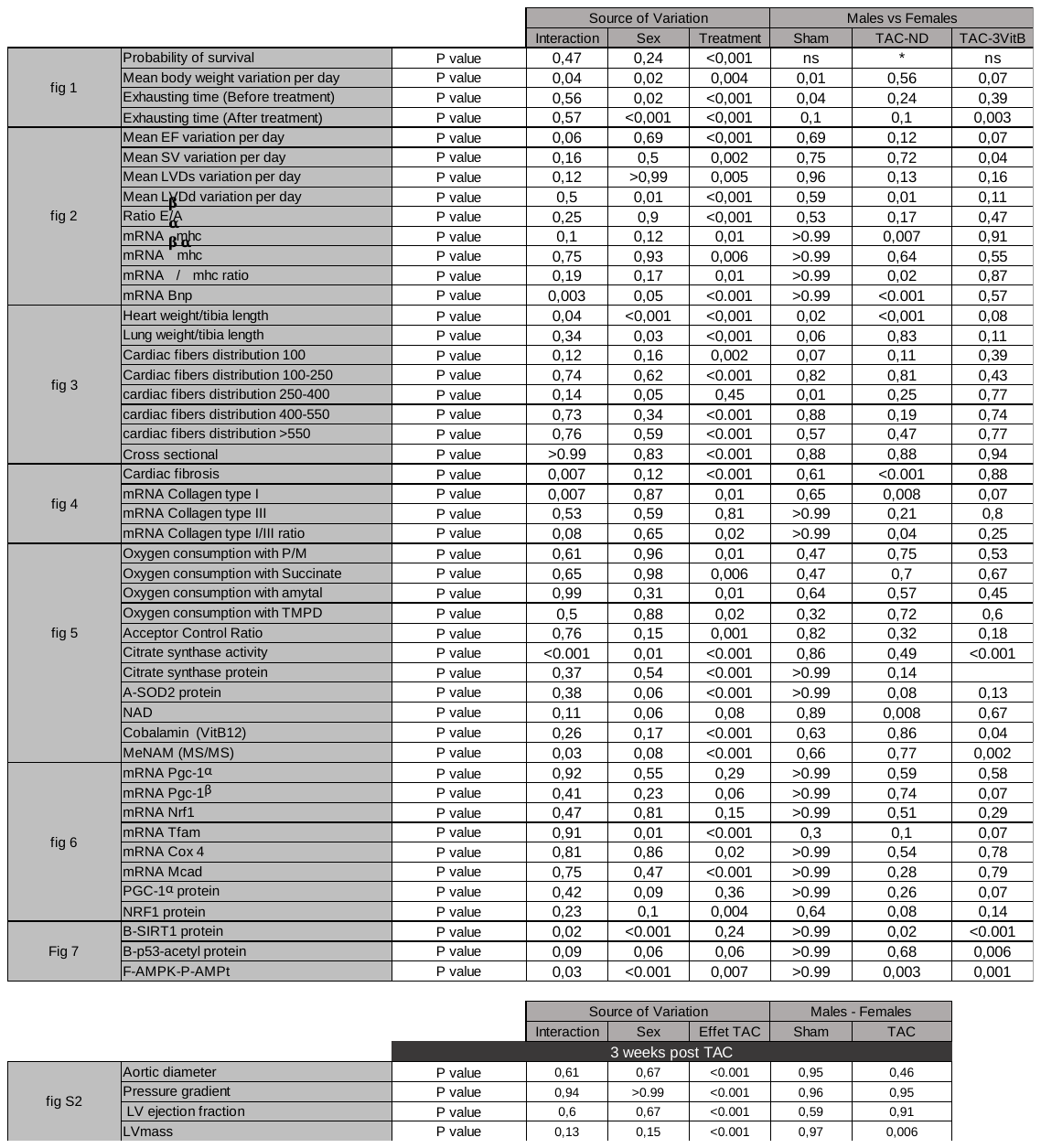

Statistical analysis.
